# Lack of Single Amino Acids Transcriptionally Remodels Sensory Systems to Enhance the Intake of Protein and Microbiota

**DOI:** 10.1101/2024.07.23.604616

**Authors:** Gili Ezra-Nevo, Sílvia F Henriques, Daniel Münch, Ana Patrícia Francisco, Célia Baltazar, Ana Paula Elias, Bart Deplancke, Carlos Ribeiro

**Author notes:** Authors contributed equally (Gili Ezra-Nevo, Sílvia F Henriques).

## Abstract

Adequate intake of dietary essential amino acids (eAAs) is vital for protein synthesis and metabolism. Any single eAA deprivation is sufficient to increase protein intake in *Drosophila melanogaster*. How such nutritional “needs” are transformed into behavioral “wants” remains poorly understood. We derived transcriptomes from the heads of flies deprived of individual eAAs to identify mechanisms by which this is achieved. We found that, while specific eAA deprivations have unique effects on gene expression, a large set of changes are shared across deprivations. We show that *Or92a* upregulation upon eAA deprivation increases the exploitation of yeast, the main protein source of flies. Furthermore, *Ir76a* upregulation was crucial for feeding on *Lactobacillus*, a gut bacterium that ameliorates the fitness of eAA-deprived flies. Our work uncovers common and unique transcriptional changes induced by individual eAA deprivations in an animal and reveals novel mechanisms underlying the organism’s behavioral and physiological response to eAA challenges.

## Introduction

Diet plays a crucial role in determining the overall health and well-being of organisms. Because they cannot be synthesized or cannot be synthesized efficiently, essential amino acids (eAAs) are crucial dietary components that play an important role in regulating the organism’s fitness. Indeed, deficiency of any eAA can be harmful for fecundity, development, growth and metabolism, energy homeostasis and brain functions from humans to flies^1–4^. It is, thus, critical that animals can detect the lack of eAAs and mount a homeostatic response to appropriately cope with such deficiencies.

The effects of nutritional state on the organism have been investigated beginning with pioneering transcriptomics work using microarrays. Focusing on the *Drosophila* larval stage, these studies characterized transcriptional changes at the whole animal level upon full starvation or yeast deprivation^5,6^. In vertebrates, dietary protein manipulations have been shown to significantly influence gene expression, leading to the identification of FGF21 as being transcriptionally responsive to dietary proteins, thereby affecting growth and metabolism^7,8^.

These and numerous follow-up nutrigenomics studies revealed profound changes in gene expression upon nutritional manipulations. Nonetheless, the extent to which these changes are driven by the lack of AAs in general and how individual AAs drive specific gene expression alterations remains poorly addressed. This is highly relevant as each AA has specific roles beyond serving as the building blocks for protein synthesis. Methionine, for example, plays a unique role in translation as the AA encoded by the start codon and serves as a precursor for methylation reactions^9–11^. Additionally, different eAAs have well-described unique impacts on health span and brain function^11–20^. Prominent examples are tryptophan from which serotonin is synthetized, and arginine which serves as a precursor for nitric oxygen production^9,21^. Indeed, several eAAs deprivations have been shown to induce gene expression changes both in cell lines and in the context of the whole animal^16,22–24^. It is thus still unclear what the core expression responses to any eAA dietary deprivation are, and to which extent specific AA deprivations lead to unique gene expression changes in the animal.

Behaviorally, deficiencies in dietary proteins or eAAs lead to a preference for protein-rich foods^25–30^. Remarkably, the removal of any of the ten eAA can be sufficient to induce a potent protein or AA appetite, as shown in *Drosophila melanogaster*^3^. Therefore, despite significant differences in eAA functions, a deficiency in any of them can induce a similar feeding-related phenotype. These feeding decisions critically rely on chemosensory input, with taste and olfaction mediating the detection of the nutritional content of food, its toxicity, and its location^31–34^. Consequently, sensory processing has been shown to be modulated by the dietary state of the animal across phyla^32,35–42^. This modulation allows animals to adapt feeding-related behaviors to their nutritional needs and achieve homeostasis. These behavioral changes have been mostly thought to rely on alterations in circuit activity at the level of sensory neurons^31,38,43–45^ or downstream sensory processing circuits^19,46–48^. However, in *Caenorhabditis elegans*, changes in chemosensory receptor expression are widely recognized as contributing to the impact of internal states on sensory processing (for example^49–51^). This stands in contrast to other animals, where less is known about the contribution of diet-induced changes in chemosensory receptor expression to maintaining nutritional homeostasis. Recent work indicating that transcriptional dynamics play a crucial role in adapting sensory sensitivity in vertebrates suggests that transcriptional changes may have a greater impact on behavioral homeostasis across phyla than previously appreciated^52,53^.

Under nutrient deprivation conditions, the gut microbiome has emerged as a crucial modulator of host physiology and behavior^54–62^. We have, for example, shown that in *Drosophila*, gut bacteria, including *Lactobacilli*, significantly impact feeding behavior and reproduction in eAA-deprived animals^3,56^. Furthermore, *Lactobacilli* have been shown to play a vital role in the growth and development of *Drosophila* larvae and juvenile mice, especially when nutritional resources are limited^63–65^. Taken in combination with the demonstrated impact of *Lactobacilli* on mouse behavior^66^ and human health^67^, the combined, available data, point to this bacterial genus as a key player in the ability of the microbiome to influence host fitness across species. Intriguingly, we and others have shown that animals can seek out food laced with gut bacteria, indicating that organisms can self-regulate the composition of their microbiome^3,68^. To what extent this is the case, what the potential mechanisms are that mediate this ability, and how it might be adapted to the current needs of the animal remains poorly understood.

In our work, we have utilized the fruit fly to generate a comprehensive overview of eAA depletions. Using synthetic diets, we have exposed flies to diets depleted of each of the 10 eAAs, followed by RNA-sequencing (RNA-seq) of heads (comprised of the brain, sensory organs and a key metabolic organ, namely, the fat body). We have generated a complete gene expression map following each eAA deprivation unraveling the unique expression “fingerprint” of each eAA depletion. We have also found that eAA depletion leads to a common change in gene expression, including in olfactory-receptor genes. We further show the relevance of the regulated receptors to feeding choices made by the fly. *Or92a* regulation is important for the flies’ increased engagement with protein-rich foods (namely, yeast) upon single eAA deprivations. *Ir76a* expression is also sensitive to amino acid levels in the food; however, it remarkably influences the flies’ approach towards a bacterium from the *Lactobacillus* genus, which is part of the natural fly microbiome. Our work maps characteristics of specific eAA depletions on the fly head transcriptome across all eAAs. We unravel a mechanism by which a dietary need induces specific changes in sensory processing that are translated into attraction towards appropriate food sources and beneficial gut bacteria.

## Results

### Dietary depletion of any essential amino acid induces a strong preference for proteinaceous food

Dietary proteins are essential components of animal diets as they are the main source of essential amino acids (eAAs). As such, animals, including *Drosophila melanogaster*, react strongly to dietary deprivations of proteins or AAs by altering their feeding preferences^3,25,31,69,70^. When given the choice to feed from either a proteinaceous or a carbohydrate food source, flies deprived of any single eAA switch their preference from sucrose towards yeast (their main dietary source of proteins) feeding ^3^. One would expect that this change in preference is mainly driven by a strong increase in the intake of proteinaceous food. Indeed, a three-day deprivation of any single eAA in mated female flies (Fig. 1A-C) led to a strong increase in total feeding on yeast compared to flies kept on a complete medium as measured using the flyPAD technology (Fig. 1B-C). The effect was nutrient-specific as no change was observed for sucrose feeding (Supplementary Fig. 1A). Classic work has shown that specific AAs, especially branch chain AAs (BCAAs) are key modulators of TOR signaling and therefore cell physiology^71,72^. Furthermore, different eAAs are represented differently in the exome^73^ and support different metabolic processes beyond translation^9^. Nevertheless, the depletion of any of the ten eAAs from the diet induces a robust and consistent increase in yeast intake, suggesting that a common mechanism could underlie this change in yeast preference.

**Fig. 1.**
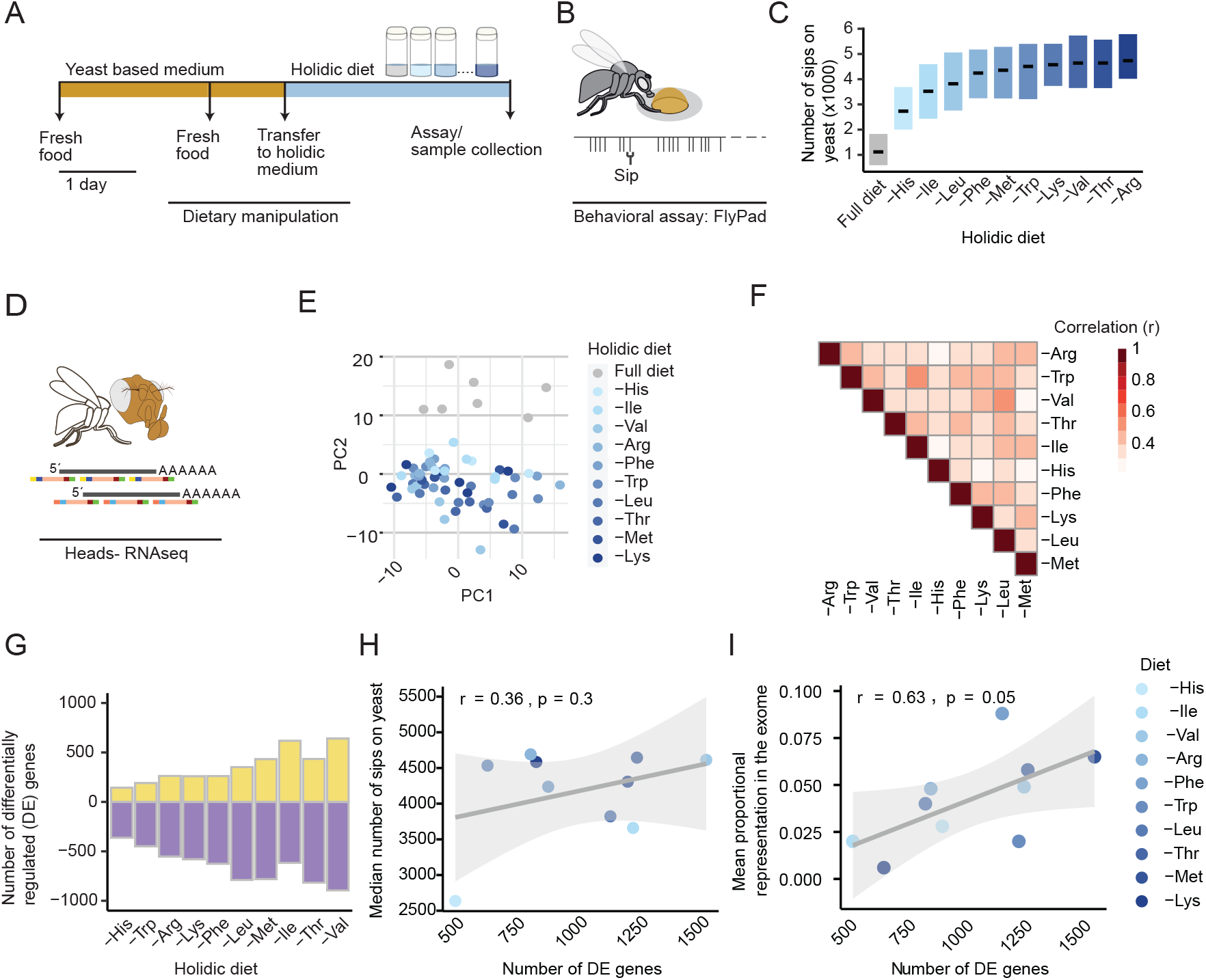
Single amino acid deprivation is sufficient to induce changes in feeding and head transcriptome **A-C:** Procedure: Flies were sorted to yeast based medium, transferred to fresh food after 2 days, and then transferred to holidic media containing all nutrients (control) or lacking a single eAA, for 3 days **(A)**, followed by behavioral assay to assess yeast feeding using the flyPAD (**B**). Black lines represent the medians and the boxes represent 95% confidence interval (CI) of the number of sips on a yeast food source measured for fully fed flies (gray) or single eAA deprived flies (shades of blue) **(C)**. From the same batch flies’ heads were dissected (heads) for RNA sequencing (RNAseq) (**D**). **(E)** Principal component plot of all samples (62) based on the top 500 most variable genes. **(F)** Heatmap depicting the Pearson correlation between each pair of deprivations. Values used are the log2(fold change) of each deprivation compared to the full diet condition for all the genes sequenced. **(G)** Number of up-(yellow) and down-regulated (purple) genes per deprivation condition. **(H)** Pearson correlation between the number of differentially expressed (DE) genes and the median number of sips on yeast as shown in (C), gray shade represents 95% CI. **(I)** Pearson correlation between the number of differentially expressed (DE) genes and their relative representation in the exome (according to Piper et al., 2017).

### Single eAA deprivations are sufficient to induce changes in expression

We hypothesized that the mechanisms driving the coherent behavioral change across all eAA deprivations as well as the unique contribution of each eAA to cellular functions could be at least in part mediated by transcriptional changes. We therefore sequenced RNA of heads of mated female flies, each deprived of a different eAA, to identify the transcriptomic changes associated with the deprivation of each of the ten eAAs (Fig. 1D). The fly head, which is a heterogeneous body part, contains key tissues related both to controlling behavior (the brain and sensory organs) and metabolism (the fat body). As such, we hypothesized that changes in gene expression altering behavior as well as metabolism, upon eAA deprivation, should be reflected within this body part. Heads from flies exposed to a diet depleted of any eAA showed a consistently different transcriptome compared to flies kept on a complete diet (Fig. 1E; total un-normalized counts also differed, see Supplementary Fig. 1B). At this analysis level, the different deprivations did not separate from each other based on the most variable genes, suggesting that the changes in gene expression can be largely explained by a common set of genes.

Nonetheless, when correlating the change in transcriptome following each eAA elimination from the diet between the different conditions (Fold Change of all genes from control), there was less correlation than expected (Fig. 1F). It is thus clear that each eAA deprivation also induces unique changes in the transcriptome. Of note, the correlation between Leu and Val, both BCAAs, is relatively high while the third BCAA, Ile, seems to be better correlated with Trp. Furthermore, no deprivation effects were anti-correlated. Counting the number of differentially expressed (DE) genes, we found that between 400 (-His) and 1600 (-Val) genes were significantly DE following each eAA deprivation with more genes being downregulated than upregulated in all conditions (Fig. 1G; Figure S1C). The exception seemed to be Ile deprivation where the number of up and down-regulated genes is roughly the same. Overall, 195 genes were significantly regulated in all 10 deprivations, and 308 in at least 9 conditions (adjusted p value <0.1).

Importantly, Leu, Met, Ile, Thr, and Val depletion induced a larger number of DE genes compared to the other eAAs (over 2-fold difference in number of DE genes; Fig. 1G; Figure S1C). Of these, Ile, Val, and Leu are BCAAs, which have been shown to have a significant impact on metabolism^4^ and are potent modulators of TOR signaling and therefore gene transcription^71,74^. Furthermore, given its unique role in translation, Met is widely appreciated for inducing potent and unique changes in cell and animal physiology^14,18^. Overall, our data suggest that dietary eAA deprivation induces significant gene expression changes. A common set of genes explains the differences between fully fed and eAA-deprived animals, while individual AAs differ in their ability to control the expression of specific genes.

### The number of DE genes following eAA deprivation is correlated to their proportion in the exome, but not behavior

We wondered if the number of DE genes induced by each deprivation is correlated with the strength of the motivation to feed on yeast. To answer this, we used the median number of sips on yeast following each eAA depletion and correlated it with the total number of DE genes upon that specific dietary intervention. The number of differentially expressed genes showed a low correlation with the behavioral parameter (Fig. 1H; r=0.36, p=0.3). This implies that it is not the number of genes being modulated (i.e. the global gene expression impact of the deprivation) but rather which genes are modulated that could explain the changes in behavior. We then tested whether the predicted representation (and therefore predicted rate of use at the translational level) of each eAA in the exome correlates with the measured impact of its depletion on the transcriptome. For this, we correlated the proportional representation of AAs in the exome as calculated by Piper and colleagues^73^ with the number of differentially expressed genes upon the corresponding eAA deprivation (Fig. 1I). There was indeed a strong correlation between the level at which a specific eAA is predicted to be used during translation, and how much its depletion impacted changes in gene expression (r=0.63, p=0.05, based on the FLYAA diet; Fig. 1I). Methionine is a unique eAA as it is the translated AA of the start codon, as such it is expected to have an effect on the synthesis of proteins that far exceeds its mere representation in the exome. We thus wondered if removing Met from the correlation analysis would reveal a stronger correlation between the 9 other eAAs and the exome, which indeed was the case (r=0.76, p=0.017; Figure S1D). Therefore, the predicted demand for eAAs during translation seems to emerge as a significant underlying factor explaining the impact of each eAA deprivation on gene expression. In contrast, the behavioral response appears uncorrelated with the number of differentially expressed genes, suggesting that the observed bulk of the changes in mRNA expression do not account for the observed changes in behavior.

### Ile and Met are unique modifiers of the transcriptome

To further explore the similarities and differences between the different eAA depletions we focused on the top 20 most significantly regulated genes in each deprivation and clustered these according to the pattern of normalized gene expression (Fig. 2A). Many of these reliably regulated genes have previously been shown to be sensitive to dietary manipulations, especially changes in dietary protein levels (Figure S2A). These include the genes encoding storage proteins (*Lsp2*, and *Yolk proteins 1-3*^75^) as well as *fit*^76^, and *CCha2*^77^(green boxes in Fig. 2A and Figure S2A^78,79^). Distance-based clustering analysis further validated not only the difference of all deprived samples from the full-diet control, but also the unique impact of two eAAs in particular, namely Ile and Met (Fig. 2A). Importantly, genes known to be transcriptionally responsive to Met deprivation such as *methionine sulfoxide reductase A (Eip71CD also known as MsrA;*^16^) were also identified as such in our dataset. Under the -Ile condition, several unique genes are regulated as well (such as *Arc1* and *eIF6*). To conclude, the removal of individual eAAs from the diet modulates a large core set of targets common to all eAA deprivations, while each eAA depletion also induces a unique transcriptional footprint.

**Fig. 2.**
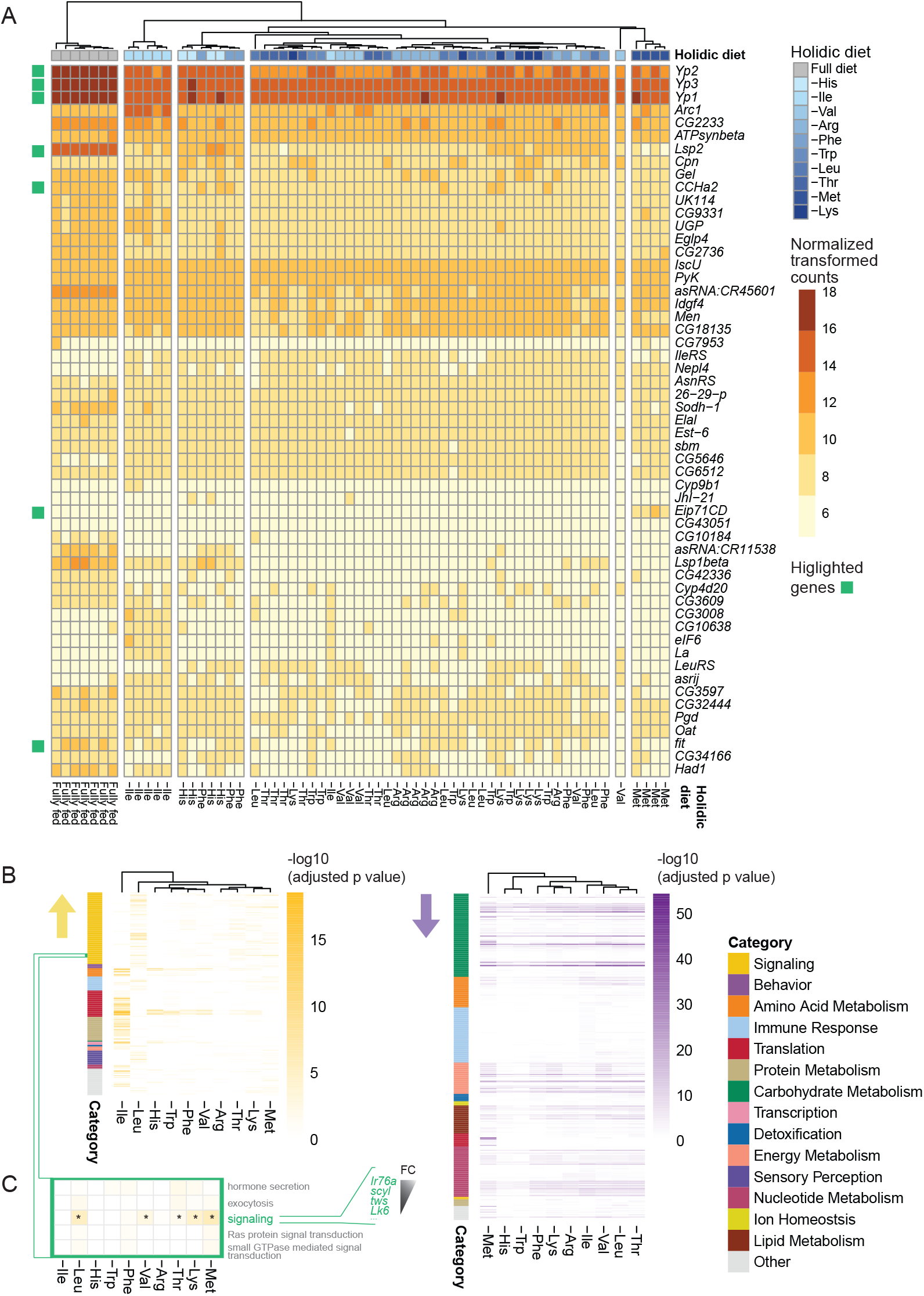
Ile and Met deprivations induce a unique transcriptional regulation fingerprint in the fly head. **(A)** Distance based hierarchal clustering based on top 20 most significant DE genes per eAA deprivation (pHeatmap.r); Shades of orange-brown represent normalized-transformed values (Deseq2.r); Green squares refer to genes highlighted in the text (**B)** Distance based hierarchal clustering using all gene ontology (GO) terms enriched in any eAA deprivation (FDR adjusted p<0.05) for upregulated genes (yellow) and downregulated genes (purple; pHeatmap.r); Colors on left side of plots represent manual categorization of GO term. **(C)** Highlighted GO terms within the signaling category; yellow shades represent -log10 of FDR adjusted p-value; stars within the signaling GO term highlight eAA deprivations for which adjusted p-value<0.05; genes depicted in green are genes within the signaling GO term that were differentially expressed in most (9 -10) dietary conditions. They are ordered according to mean fold changes (FC).

### Similarities across eAA deprivations are more pronounced in downregulated pathways, while upregulated pathways vary

To identify the molecular processes modified under the different eAA deprivations, we used gene ontology (GO) analysis based on either upregulated or down-regulated genes (adjusted p-value <0.1). We furthermore focused on the GO terms that were enriched following any deprivation (i.e. significant for at least one dietary condition; Fig. 2B) and the top 5 most significant terms per diet (Figure S2B). This revealed that while upregulated pathways were quite varied across the different eAA deprivations (Fig. 2B, yellow; Figure S2B, yellow), most downregulated pathways were found to be generally common across the deprivations (Fig. 2B, purple; Figure S2B, purple). Overall, most changes were related to processes relevant for managing the cellular impact of a decrease in translation. Downregulated genes were enriched for metabolic and translation processes which indicates that the animal reduces bulk translation and energy use (including the production of storage proteins) to account for a global reduction in protein synthesis (Fig. 2A and B; Figure S2B). To counter the shortage in available AAs, cells seem to also increase the expression of genes encoding for proteins responsible for enhancing the bioavailability of AA for protein synthesis. These are for example tRNA synthetases which are upregulated upon eAA deprivation (*Il-eRS, AsnRS, LeuRS*). Overall, the bulk of the changes in expression observed upon eAA deprivation can therefore be attributed to the attempt of the animal to counter a shortage in eAA available for protein translation.

Distance-based clustering of the different GO terms enriched in any of the AA deprivations highlighted the unique signature of Ile and Leu deprivations, which was mainly visible in the upregulated category, similar to what was observed in vertebrate studies (Fig. 2B, yellow)^80,81^. In the downregulated pathways, again, Met showed a distinct signature in the translation category (Fig. 2B, purple; Figure S2B). This highlights that specific eAA deprivations can also regulate specific biological processes, suggesting that animals can sense the bioavailability of specific AAs. How the identity of specific AAs is sensed and how this is translated into a specific biological output are poorly understood phenomenona.

### Single eAA depletions change the expression level of specific odorant receptors

One of the categories that showed the most consistent upregulation across all eAA deprivations was the signaling category (Fig. 2C). The highest fold change within that category was observed for *Ir76a*, which encodes an olfactory receptor^82,83^. This led us to test if other olfactory receptor genes were regulated to support the flies’ foraging towards the appropriate food sources. We thus focused on analyzing changes in the expression of chemosensory genes (Fig. 3A). This analysis revealed two genes, encoding olfactory receptors, *Ir76a* and *Or92a*, as being consistently upregulated in all deprivation conditions (Fig. 3A). We next validated the upregulation of *Or92a* using qPCR, confirming that the deprivation of a single eAA, namely Ile, was sufficient to induce the predicted increase in expression observed upon eAA deprivations (Fig. 3B). To validate this result further, we used a transgenic line as a reporter for the regulation of the olfactory receptor gene expression (*Or92a-Gal4*). Indeed, eAA deprivation led to an increase in reporter expression levels (Fig. 3C). Taken together, these results indicate that eAAs modulate the olfactory receptor expression repertoire of flies and that *Or92a* is upregulated upon eAA depletion.

**Fig. 3.**
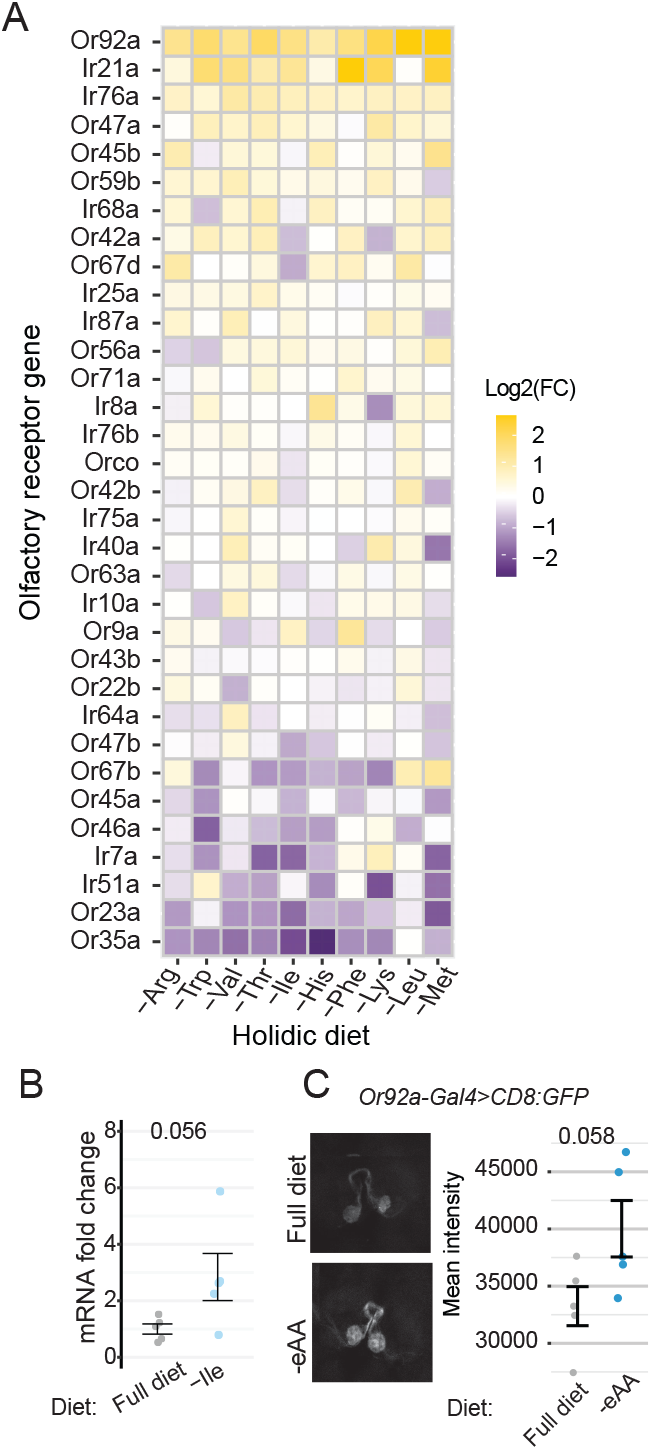
Specific olfactory receptor genes are transcriptionally upregulated by any single eAA deprivation **(A)** Heatmap depicting log2 fold changes (FC) of olfactory genes expression following each eAA deprivation compared to control flies on full diet. **(B)** Validation of *Or92a* mRNA regulation by diet. *Or92a* mRNA level fold change in heads obtained from flies following 3 days on diet deficient in Ile compared to control (full) diet; data presented as +-standard error of the mean (SEM), each data point represents a pooled sample of 50 fly-heads; P-values are obtained by performing Wilcoxon signed rank test, and are indicated at the top of the plot; n=5; red squares indicate number of conditions an Or/Ir was significantly DE (p<0.1) **(C)** Reporter based validation of *Or92a* regulation by diet. Left panel: representative sum of projections confocal images of antennal lobes obtained from *Or92a-Gal4> CD8:UAS-GFP* flies following 3 days on synthetic full diet or diet lacking all eAAs. Right panel: Quantification of normalized mean fluorescence intensity in *Or92a* glomeruli; data presented as +-SEM. Each data point corresponds to an individual fly; P-values are obtained by performing student-t test, and are indicated at the top of the plot. n=5.

### eAA depletion-induced *Or92a* upregulation is important to sustain flies’ engagement with yeast

Dietinduced changes in olfactory receptor expression are likely to modulate how the fly finds and interacts with food. Indeed, protein- and AA-deprived flies remodel their foraging behavior to increase the efficiency by which they engage with yeast patches, at the expense of exploration^43,84–86^. To dissect the functional relevance of the observed change in olfactory receptor expression, we therefore chose to use a behavioral tracking assay in which flies forage in an arena containing yeast and sucrose food patches (Fig. 4A-B)^84^. First, we validated that a single dietary eAA depletion (Ile) is sufficient to induce a similar behavioral effect as we had previously observed upon full AA deprivation (Fig. 4C-D)^84,86^. We focused on the overall exploration of the arena, represented as the number of encounters (number of times a fly encountered the food spot), the number of visits to a food spot (the number of times a fly stopped at a food spot and started feeding). And the exploitation of the food, represented by the amount of time flies spent on a food spot (represented by two metrics: the total time on a food spot during the 1h assay and the length of the longest visit to a food spot) (Fig. 4B). Flies deprived of a single eAA (blue) increased their interaction with yeast (total time spent on a food patch and longest visits increased compared to control, i.e. fully fed flies (grey); Fig. 4D). Ile-deprived flies also increased their visit number to yeast spots but reduced overall exploration (the number of encounters went down as flies spent more time on food spots).

**Fig. 4.**
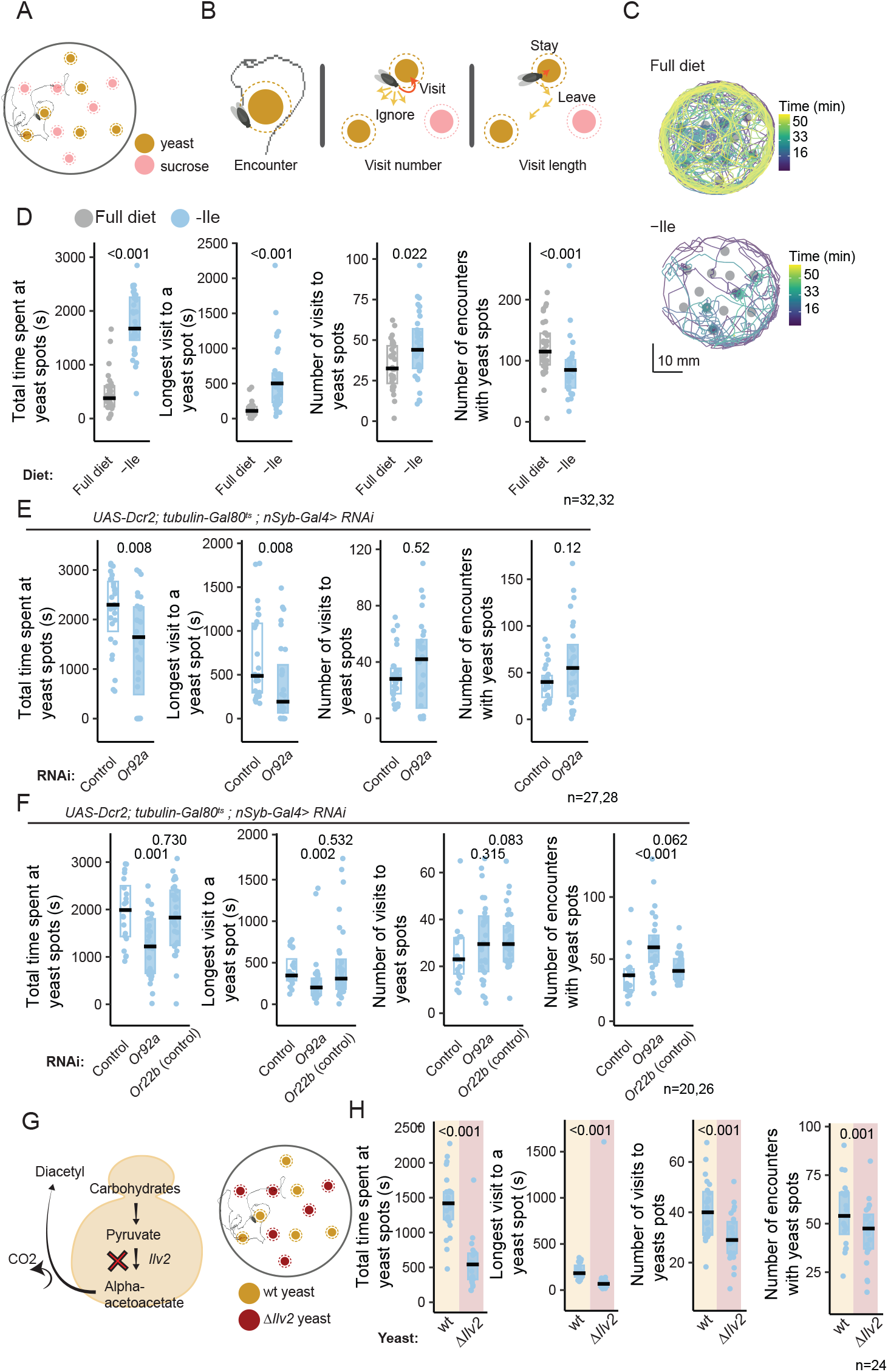
Internal state driven upregulation of *Or92a* drives engagement with yeast **(A)** Layout of the tracking arena. Each arena holds a single fly, six yeast and six sucrose spots. **(B)** Behavioral metrics generated using the tracking assay: number of times a fly passed on the yeast spot (encounters), number of times a fly stops and feeds at a yeast spots (visits), and the duration of each visit (visit length). Total time spent on a food spot = mean duration * number of visits. **(C)** Representative trajectories of a flies foraging for 1 hour in an arena containing sucrose and yeast patches following 3 days on a full diet or on diet lacking Ile. Color scale represents time from beginning of the assay. **(D)** Foraging behavior values as read out using the tracking assay. Behavior of control flies maintained on either a full diet (open-gray boxes) or deprived of Ile (full light blue boxes). Here and in other behavior plots, the black line represents medians, boxes represent inter-quartile range (IQR). Each datapoint corresponds to a single fly. Numbers at the top of the plots indicate p values as calculated using the Wilcoxon signed rank test. n=32. **(E)** Tracking assay results of Ile deprived flies expressing RNAi (time restricted) against *Or92a* (full light blue boxes) or control RNAi (open light blue boxes) driven by *UAS-Dcr2; tubulin-Gal80*^*ts*^; *nSyb-Gal4*. Numbers at the top of the plots indicate p values as calculated using the Wilcoxon signed rank test; n=27,28. **(F)** Tracking assay results of Ile deprived flies expressing RNAi against *Or92a* (full light blue boxes), control RNAi (open light blue boxes) or against *Or22a*, used as control olfactory receptor target (open light blue boxes); Genotypes are indicated at the top of all plots. n=20-32. **(G)** Schematic representation of yeast with *Ilv2* mutation (*Ilv2*; left) and of the arena setup for the choice between *wt* yeast (results on brown background) and *Ilv2* yeast (results on dark red background). **(H)** Tracking assay results comparing wt flies’ behavior towards *wt* yeast (brown) and *Ilv2* yeast (Burgundy red) in the same arena. n=24.

Since odor could play an important role in yeast foraging, we hypothesized that the transcriptionally modulated odorant receptors could play a role in the detection or attraction to the food patches.

To validate the specific effect of receptor upregulation upon AA deprivation, we decided to test Ile-deprived flies in which we knocked down *Or92a* receptor (thus reversing the effect of diet-induced upregulation of the receptor). Ile-deprived flies, in which we acutely thermogenetically knocked down *Or92a* showed reduced engagement with yeast patches compared with Ile-deprived control flies (as measured using the total time spent on the food patch during the assay and the longest time spent on a patch during one visit; Fig. 4E). *Or92a*-expressing olfactory neurons have been suggested to play an important role in directing the fly towards vinegar, an important product of fruit fermentation^87–89^. We therefore tested if the reduction in total engagement with yeast (the source of fermentation) was due to a decrease in the ability of the fly to find the food. Interestingly, the number of times the fly engaged with the yeast spot during the assay was not affected by *Or92a* knockdown (as measured by the number of visits; Fig. 4E). Importantly, the number of times a fly encountered a food patch (with and without stopping to feed, indicated by the number of encounters) was also not decreased, indicating that the motivation or ability of the fly to explore the arena was not affected. *To confirm that not any modulation of a single olfactory receptor impacts the fly’s food-foraging parameters we knocked down another olfactory receptor gene for which we observed no change in our RNA-seq data* (*Or22b). Indeed, Or22b knockdown* did not show a phenotype when compared to controls, indicating the observed phenotypes were specific for *Or92a* (Fig. 4F). Thus, the reduced duration of engagement with the yeast spots cannot be explained by lack of exploration, immobility, or ability to find yeast but suggests that Or92a modulates the way eAA-deprived flies exploit yeast patches once they find them.

### Production of diacetyl, a key ligand for Or92a, is crucial for flies’ engagement with yeast

*Or92a*-expressing sensilla have been shown to mediate the detection of diacetyl (2,3-butanedione) by the olfactory system^90–92^. Based on our data, one would expect that the production of diacetyl by yeast would be key for mediating how the fly exploits yeast. In yeast, *Ilv2* encodes an enzyme responsible for the synthesis of diacetyl (Fig. 4G). We, therefore, decided to compare how flies forage for control yeast and yeast that are mutants for the *Ilv2* gene and thus have impaired diacetyl production^93,94^. Indeed, eAA-deprived flies showed a clear preference for control yeast over yeast with the mutation in the diacetyl production pathway, indicated by the shorter total time spent on the mutant yeast and shorter visit length (Fig. 4H). Importantly, this phenocopies the effect of knocking down *Or92a* in the fly (Fig. 4E-F). However, flies also showed fewer encounters and fewer visits to the mutant yeast (Fig. 4H), implying that other receptors detect diacetyl in yeast as well. These receptors would be important for the fly to find yeast, which in our assay is likely to be independent of Or92a. To conclude, we have shown that *Or92a* is important for flies to efficiently exploit yeast and that the detection of yeast-derived diacetyl odor by eAA-deprived animals is likely to be a key process that increases the engagement of flies with yeast upon eAA deprivation. This modulation of OR expression therefore contributes to the ability of eAA-deprived flies to remodel their behavior to efficiently achieve protein homeostasis.

### Ir76a plays a key role in detecting and ingesting specific gut bacteria

Another olfactory receptor gene that was consistently and significantly modulated by eAA deprivation was *Ir76a* (Fig. 3A). Similar to what we did for *Or92a*, we validated our finding using a transgenic reporter (*Ir76a-Gal4*) and observed a significant increase in the fluorescent signal under eAA deprivation conditions (Fig. 5A), something we did not observe for another olfactory receptor gene for which we observed no change in our RNA-seq data (*Ir64a; Fig. 5B*). To test if high levels of dietary proteins suppress *Ir76a* expression, we increased the dietary yeast available to the fly for 6 days. This led to a strong and significant decrease in the signal of the expression reporter for *Ir76a* (Fig. 5C), and remarkably only 3 days on high protein was sufficient to induce a decrease in *Ir76a* reporter expression, albeit to a smaller extent (Figure S3A).

**Fig. 5.**
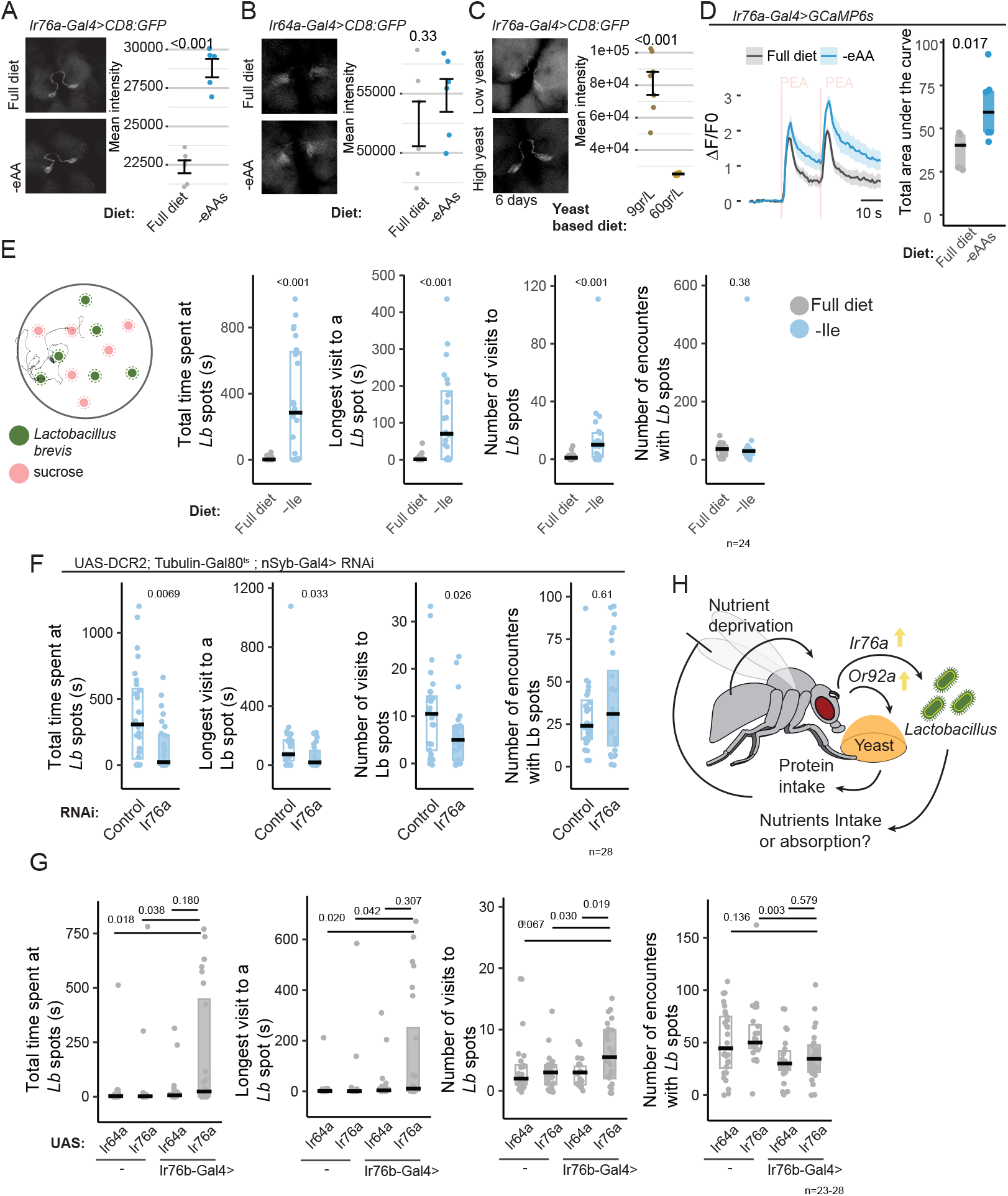
Nutritional state dependent *Ir76a* upregulation drives attraction towards gut bacteria **(A)** Reporter based validation of *Ir76a* regulation by diet. Left panel: representative sum of projections confocal images of antennal lobe obtained from *Ir76a-Gal4> CD8:UAS-GFP* flies following 3 days on synthetic full diet or diet lacking all eAAs. Right panel: quantification of normalized mean fluorescence intensity in *Ir76a* glomeruli; data presented as +-SEM. Numbers at the top of the plots indicate p values as obtained using a t-test; n=5. **(B)** Reporter based quantification of *Ir64a* regulation by diet. Left panel: representative confocal images of antennal lobes obtained from *Ir64a-Gal4> CD8:UAS-GFP* flies following 3 days on synthetic full diet or diet lacking all eAAs. Right panel: quantification of normalized mean fluorescence intensity in *Ir64a* glomeruli. Data presented as +-SEM. Numbers at the top of the plots indicate p values as calculated using a t-test; n=6. **(C)** Reporter based analysis of *Ir76a* regulation by yeast content in the diet. Left panel: representative confocal images of antennal lobe obtained from *Ir76a-Gal4> CD8:UAS-GFP* flies following 6 days on standard lab food containing 9gr yeast/L or 60gr yeast/L. Right panel: quantification of normalized mean fluorescence intensity in *Ir76a* glomeruli. Data presented as +-SEM. Numbers at the top of the plots indicate p values as calculated using a t-test. n=7. **(D)** Left: mean Phenylethylamine (PEA; red vertical lines represent the stimulus duration) response traces from *Ir76a* glomeruli in 5 flies on full diet (gray) or 6 on a diet lacking eAAs (dark blue); Shading represents +-SEM. Right: Box plot comparing response magnitudes of response to PEA in *Ir76a* glomeruli in flies on full diet (gray) or eAA deprived (dark blue). Numbers at the top of the plots indicate p values as calculated using the Wilcoxon signed rank test. n=5-6. **(E)** Left: schematic representation of the tracking assay layout with six patches containing *Lactobacillus brevis* (*Lb*) and six containing of sucrose. Right: *Lb*-tracking assay results comparing flies maintained on full diet (gray boxes) or a diet lacking Ile (light blue boxed). Black line represents median values. Boxes represent IQR. Each datapoint corresponds to a single fly. Numbers at the top of the plots indicate p values as calculated using the Wilcoxon signed rank test. n=24. **(F)** *Lb*-tracking assay results of Ile deprived flies expressing RNAi against *Ir76a* (full light blue) or control RNAi (open light blue) driven by *UAS-Dcr2; tubulin-Gal80*^*ts*^; *nSyb-Gal4*. Driver line indicated at the top of all plots. Black line represents medians. Boxes represent IQR. Each datapoint corresponds to a single fly. Numbers at the top of the plots indicate p values as calculated using the Wilcoxon signed rank test. n=28. **(G)** *Lb*-tracking assay results of fully fed flies with *Ir76a* or *Ir64a* (control ionotropic receptor) ectopic expression driven by *Ir76b-Gal4*, and UAS-insertion site controls. Black line represents median values. Boxes represent IQR. Each datapoint represents a single fly. Numbers at the top of the plots indicate p values as calculated using the Wilcoxon signed rank test. n=23-28. **(H)** Proposed model for olfactory receptor modulations under dietary perturbations and their relevance to regaining homeostasis.

Ir76a was shown to respond to Phenylethylamine (PEA)^83,95^. Thus, to validate the functional relevance of *Ir76a* upregulation under eAA depletion conditions, we tested whether the response of *Ir76a*-expressing sensory neurons to PEA was modulated by eAA deprivation. Indeed, Ca^2+^ imaging of these neurons at the level *of the* glomeruli showed greater response to PEA in deprived flies compared to fully fed flies, providing functional support for the relevance of the observed diet-induced upregulation of *Ir76a* (Fig. 5D).

### Ir76a is important for the attraction to gut bacteria under deprivation conditions

Intriguingly, PEA has been reported to be the product of AA fermentation by *Lactobacillus brevis (Lb)*^96^. We previously demonstrated that *Lb* is a key component of the fly microbiome due to its ability to regulate protein appetite and improve egg production in AA-deprived females^3,56^. Furthermore, it has been shown that in *Drosophila* larvae, *Lactobacilli* can improve metabolism and nutrient absorption from the food^54^ and that flies have a preference for food laced with gut bacteria^3,68^. *We therefore asked whether Lb is attractive to flies under deprivation conditions and whether Ir76a could play a role in this attraction*. We modified our tracking assay to give Ile-deprived flies a choice between *Lb* and sucrose and tested the effect of acutely knocking down *Ir76a* on this choice (Fig. 5E-F). In agreement with the Ca^2+^ imaging data, we found that in control flies, Ile deprivation dramatically increases the attraction of flies towards *Lb* (Fig. 5E). Furthermore, thermogenetically-induced knockdown of *Ir76a* inhibited this preference (Fig. 5F). More specifically, *Ir76a* knock-down flies showed a reduced number of visits to the bacteria, which is likely the main contributor to the reduced total time spent on the *Lb* spots, as we did not observe a change in the mean visit duration (Figure S3B). Furthermore, we did not observe a phenotype in fully fed flies, which is consistent with the low expression of the receptor in these animals (Figure S3C). To test whether increased *Ir76a* expression is sufficient to increase approaches towards *Lb in fully fed flies*, we over-expressed *Ir76a* in chemosensory neurons using *Ir76b-Gal4*^82^. In agreement with the changes observed upon eAA deprivation, even in fully-fed flies, the overexpression of *Ir76a* increased the approach behavior towards bacteria, as measured by the number of visits, despite their low motivation to eat. *Ir76a* overexpression flies tended to have longer visits to *Lb* spots as well, as measured by the length of the longest visit to *Lb* spots, however, again, with a smaller effect compared to the number of visits (Fig. 5G). The number of encounters with *Lb* spots remained unaffected by *Ir76a* manipulation (Fig.5G). Taken together, these results strongly suggest that *Ir76a* plays an important role in facilitating the identification of food containing a *Lactobacillus* and therefore its ingestion. Importantly, this attraction is strongly increased by eAA deprivation, likely through the increase in expression of this specific chemoreceptor. This fits with the previously described beneficial role of *Lactobacilli* in helping the fly cope with the deleterious effects of low protein and eAA diets^3,56,63,64^.

## Discussion

Since proteins are the key nutrient source for essential AAs, maintaining both the quality and quantity of protein intake is key for animal well-being^97,98^. Proteins themselves are highly heterogeneous, so the amounts of AAs obtained from them can vary. Furthermore, some eAAs have functions beyond protein synthesis as they are precursors for other metabolites, signaling molecules, and neuromodulators, making each eAA unique in its impact and importance for cellular and organismal functions. Despite efforts to understand the role and impact of single AAs on cells and organisms^18,23,71,99^, there have been very few comprehensive studies assessing the importance of all eAAs in comparable conditions in animals. Here, we performed a comprehensive analysis of the impact of removing each eAA one by one on the gene expression repertoire in fly heads, resulting in a unique dataset that enables a thorough exploration of the impact of each eAA deprivation in this animal tissue. On the one hand, this dataset reveals the general strategies organisms use to cope with overall eAA deprivation, on the other hand, it uncovers the specific effects driven by individual eAAs. Although many of these eAAs have already been described as having specific effects in the literature, the unique signatures we describe for eAAs, such as Ile and Met, highlight specific processes controlled by these AAs^15,80,81,100^. Moreover, much remains to be learned about the less-explored eAAs and their unique effects (for example His, Phe). Having identified the molecular impacts triggered by their deprivation will guide future research toward an integrated understanding of how they differentially affect behavioral or metabolic processes in animals.

At the molecular level, much is known about nutrient-sensing pathways detecting the lack of AAs in cells. While TOR signaling is thought to mainly react to the lack of specific BCAAs^101^, the GCN2 pathway is supposed to mediate the generalized detection of the lack of AAs (through the recognition of uncharged tRNAs)^102^. While these pathways have been shown to play an important role in mediating the effect of AA deprivation on aging and brain function^103,104^ there is a significant gap between our understanding of how these pathways operate in cells versus complex multicellular organisms. Importantly, TOR and GCN2 pathways are supposed to act on a common set of target genes which are independent of the specific nature of the lacking AA. Our current knowledge of these pathways can therefore not fully account for the unique transcriptional fingerprints observed for each eAA depletion. Thus, our work offers an attractive dataset to explore novel mechanisms by which the lack of individual eAAs can alter gene expression in animals. To further understand the mechanisms mediating these transcriptional regulations, it would be advantageous to assess how the mRNA landscape under single eAA depletion conditions corresponds to the regulons of various transcription factors (TFs) in specific cells. An interesting starting point to identify the mechanisms by which the AA state controls chemosensory genes might come from pioneering studies that identified many TFs controlling the establishment and maintenance of *Or92a* expression during development^105–107^. However, if and how these TFs might mediate the plasticity of expression in different nutrient conditions we describe here remains to be tested.

Importantly, animals must integrate signals about their current internal (AA) state and that of the food that is currently available. The first includes the evaluation of the current long-term nutritional state and should rely on some internal metabolic signal. The second is likely based on interaction with the “outside world” through the use of sensory cues such as smell or taste which are detected by sensory cells located on the head, appendages, or the gut of the animal^97^. Our work contributes to the evolving understanding of how external and internal sensing are integrated^36,43,108,109^. The mechanisms underlying this integration have been mainly thought to rely on the neuromodulation of circuit activity at the level of sensory neurons or downstream sensory processing circuits. Here we show that the internal metabolic state of the animal affects the expression of specific chemosensory receptors on sensory cells. Although chemosensory receptor gene expression can be found to be changed upon starvation in several studies, these are rarely highlighted by the authors or followed up functionally. These mechanisms are likely to act in parallel to other mechanisms such as synaptic facilitation in olfactory neurons upon starvation^43^. Together, they potentiate odor and taste detection leading to an increase in the attractiveness of foods that contain the nutrients required to achieve nutrient homeostasis^108^. Since chemosensation is likely to be the most ancient sense and evolved long before neurons, it is attractive to speculate that sensory modulation by regulation of chemosensory receptor gene expression might be a very ancestral form of sensory modulation.

Although odorants are very well-accepted contributors to the attractiveness of food and the ability of animals to find it, the extent to which odor perception is modulated to direct feeding-related behaviors remains poorly understood. Here, we show that the mere knockdown of a single odorant receptor gene (i.e. *Or92a*) is sufficient to disrupt the fly’s ability to reach protein-feeding levels equivalent to the controls. Importantly, this is not due to the inability of the fly to find the food, as in our experimental design, food sources are readily and abundantly available. By combining a complex foraging design with naturalistic food sources and quantitative behavioral tracking, we were able to dissect the contributions of the identified odorant receptors and their regulation in the ability of the animal to achieve homeostasis. Our work, thus, adds to previous findings showing that *Or92a* neurons are crucial for mediating the attraction to food odors^88,89,110^. Intriguingly, we find that the function of Or92a is likely to go beyond allowing the fly to find yeast patches; it also plays a role in the motivation of the fly to feed from it. Remarkably, the Solomon’s lily flower utilizes this feature to attract drosophilids by omitting yeast odors, including ligands of *Or92a*^87,111^.

*We show that Ir76a* expression is very strongly modified by the protein and AA content of the diet. Interestingly, the main function of *Ir76a* seems to be to increase the attraction of flies for *Lactobacilli*. Intriguingly, many of the ionotropic odorant receptors remain orphan receptors without known ligands or biological function. Our work suggests that they might serve as mediators between the microbial world and insects. Also, PEA is the volatile product of phenylalanine, and many volatiles made by bacteria are derived from eAAs. These volatiles might therefore serve to guide insects towards foods containing important nutrients^111^. Beyond being a source of biomass themselves gut bacteria have been shown to reprogram animal physiology and behavior. They for example increase nutrient absorption and egg production in AA-deprived animals^54–56,61,63,112^. As such, they play an important role in helping animals to cope with AA deprivation states. It is thus fascinating that AA deprivation tunes the sensory system of flies to improve their ability to find and ingest bacteria which are beneficial in this specific nutrient state. Similar mechanisms might underlie the attraction of animals towards food containing probiotics such as fermented foods. Nutrient states therefore transcriptionally reprogram chemosensory systems to fine-tune how animals interact with their microbial environment. This allows them to find and ingest both dietary sources for the nutrients they lack as well as the commensal bacteria that allow them to metabolically and physiologically optimize their physiology to better cope with the challenges of nutrient deprivations.

## ACKNOWLEDGEMENTS

We thank M. Alenius, R. C. Figueiredo, M. F. V. Dias, Z. Carvalho-Santos and the whole Behaviour and Metabolism laboratory for many fruitful discussions, valuable feedback throughout the project, and comments on the manuscript. We thanks A. Laborde for technical assistance with building the PC infrastructure for the tracking data acquisition; the Champalimaud Fly and Rodent Platforms; the Champalimaud Hardware and Software Platform; the Champalimaud Glass Wash and Media Preparation Platform for help with media preparations; we thank Lucia Serra for help with media preparation for RNA-seq behavior and sample collection experiments. We thank Joseph Paton and Georg Raiser for providing the odor stimulation device and V. Gardeux, D. Alpern and M. Demir for help with RNA—seq execution and analysis. Lines obtained from the Bloomington Drosophila Stock Center (NIH P40OD018537) and the VDRC were used in this study. G. E. N. was supported by a Marie Skłodowska-Curie postdoctoral fellowship (H2020-WF-01-2021-905234 to Gili Ezra-Nevo). APF was supported by a doctoral fellowship PD/BD/114277/2016 from the Portuguese Foundation for Science and Technology (FCT). The project leading to these results has received funding from FCT (FCT-PTDC/BIA-MOL/30081/2017), and the “la Caixa” Banking Foundation under the project codes HR17-00539 and HR23-00516 to CR.

Research at the Centre for the Unknown is supported by the Champalimaud Foundation and by Portuguese national funds through FCT - Fundação para a Ciência e a Tecnologia - in the context of the project UIDB/04443/2020, the research infrastructure CONGENTO, co-financed by Lisboa Regional Operational Programme (Lisboa2020), under the PORTUGAL 2020 Partnership Agreement, through the European Regional Development Fund (ERDF) and FCT under the project LISBOA-01-0145-FEDER-022170, and the PPBI - LISBOA-01-0145-FEDER-022122.

## Material & Methods

### *Drosophila* rearing, media, and dietary treatments

Experiments were performed with either axenic or conventional mated *w*^*1118*^ female flies unless otherwise stated. Flies were reared under controlled conditions at 25°C, 70% humidity, and 12 h light/dark cycle. Axenic *w*^*1118*^ were generated and maintained as previously described^56^ see also http://dx.doi.org/10.17504/protocols.io.hebb3an). Conventionally reared flies were maintained on yeast-based medium (YBM) (per liter of water: 8 g agar [NZYTech, PT], 80 g barley malt syrup [Próvida, PT], 22 g sugar beet syrup [Grafschafter, DE], 80 g corn flour [Próvida, PT], 10 g soya flour [A. Centazi, PT], 18 g instant yeast [Saf-instant, Lesaffre], 8 ml propionic acid [Argos], and 12 ml nipagin [Tegospet, Dutscher, UK] [15 % in 96 % ethanol] supplemented with instant yeast granules on the surface [Saf-instant, Lesaffre]). Unless otherwise stated, for dietary manipulations, Holidic medium (HM) was prepared using the exome match with optimized AA formulation (FLYAA) with food preservatives as previously described^3,113^; see also http://dx.doi.org/10.17504/protocols.io.heub3ew). HM lacking single eAAs was prepared by simply omitting the corresponding AAs. To avoid the confounding effect of nitrogen deprivation, diets lacking all eAAs (-eAA; Fig. 4 and 6) were compensated by increasing non-essential AAs quantities. For all experiments using HM the following protocol was used: 16 females and 5 males were placed in YBM-containing vials, transferred to fresh YBM after 48 h and transferred, after 24 h, to the different HM where they were maintained for 72 h before any indicated assay (Fig. 1A). For experiments in which we performed inducible knock-downs of receptors, tubes were always kept at 18°C until flies were transferred to fresh YBM, and then transferred to 29°C (4 days before behavioral assay). For experiments with increased yeast content, flies were directly sorted to YBM with either 9g or 60g yeast and without yeast granules, and transferred to similar fresh food after 3 days (a total of 6 days on YBM).

### *Drosophila* stocks and genetics

For olfactory receptors knockdown experiments, the following VDRC RNAi lines were used: #105553 (*Or92a*) and #101590 (*Ir76a*) and lines #105600 (*Or22b*) and 106267 (non-neuronal control) or 60103 (*eGFP*), crossed with *w*^*-*^, *UAS-Dcr2; tub-Gal80*^*ts*^; *nSyb-Gal4* virgin females, unless otherwise stated. Overexpression was achieved by crossing BL#41738 (*UAS-Ir76a*) and BL#41746 (*UAS-Ir64a*) with BL#41730 (*Ir76b-Gal4*). For imaging Bloomington lines #23140 (*Or92a-Gal4*), #41735 (*Ir76a-Gal4*), and #41732 (*Ir64a-Gal4*) were crossed with *UAS-CD8:GFP/CyO* virgin females.

### Sample collection for RNA-seq

For the RNA-seq dataset samples were collected in seven independent runs. Each run included control conditions (flies on a full diet) and all or most of the eAA depletion conditions, prepared as described above. From each dietary replicate, ten flies were used for tissue collection and mRNA extraction, and 16-30 flies were used to validate using the flyPADs to certify that the dietary manipulation succeeded. This was assessed by surveying that a protein appetite was induced by the deprivations.

### Total mRNA extraction, RT-PCR, and quantitative real– time PCR

For sequencing and qPCR, 10 or 50 heads, respectively, were collected from mated axenic females maintained on each dietary condition. Samples were homogenized for 30s in 350µl of PureZOL (#7326890, Bio-Rad) and RNA was extracted according to the manufacturer’s instructions (Direct-zol RNA MicroPrep #R2062, Zymo research). Samples were eluted in 50 µl of distilled RNase/DNase-free water.

### Qpcr

RNA was reverse transcribed (RT) using the iScript Reverse Transcription Supermix for RT-PCR kit (#1708841, Bio-Rad), following the manufacturer’s instructions. cDNA sample was amplified using SsoFast EvaGreen Supermix (#1725202, Bio-Rad) on the StepOnePlus™ Real-Time PCR System (Applied Biosystems) according to the manufacturer’s recommendations. The cycle program consisted of enzyme activation at 95°C for 3 s, 40 cycles of denaturation at 95°C for 5 s, and annealing and extension for 5 s. Appropriate non-template controls were included in each 96-well PCR reaction, and dissociation analysis was performed at the end of each run to confirm the specificity of the reaction. Relative RNA quantities were calculated from a standard curve and normalized to the two internal controls (*Actin42A* and *RpL32*). The primers used in this reaction were:

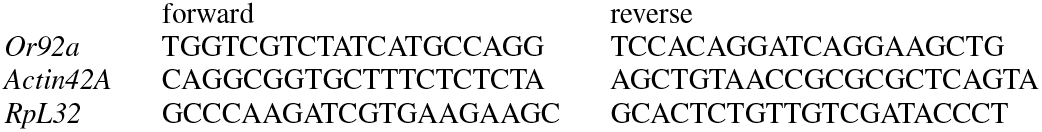

### Generation of RNAseq data and analysis

Transcriptomics analysis was performed using bulk RNA barcoding and sequencing^114^ (BRB-seq). RNA samples were barcoded and sequenced as previously described^114^. Briefly, RNA was reverse-transcribed with barcoded oligo-dT to obtain barcoded cDNA. Barcoded cDNA-RNA hybrids were pooled in a single tube and purified. The following steps, second strand synthesis, tagmentation and library indexing and amplification were all done on pooled samples. Sequencing was performed on the Illumina NextSeq 500 platform using a custom ReadOne primer (IDT) and the High Output v2 kit (75cycles) (Illumina, #FC-404-2005). Demultiplexing and alignment were done as described in Alpern et al., 2019, using genome assembly version: drosophila_melanogaster BDGP6.28 (dm6) (Ensembl release 100). Samples processing and alignment were performed in 2021 at Alithea© using their MERCURIUS service for bulk RNA barcoding and sequencing.

Counts normalization and analysis of Differentially expressed (DE) genes were done using the DESEq2 package for R with diet as the main variable and experiment date and RNA extraction date as co-variants to control for batch effect. Size factors were calculated after excluding all DE genes^115^. PCA plots were done on normalized counts after variance stabilizing transformation (vst) with DESEq2.R. DE genes for each eAA-depleted dietary condition compared to control (full diet) were determined using the cutoff of <0.1 for the adjusted p-value.

Top DE genes-based clustering: clustering analysis was done using the 20 DE genes with the lowest adj. p values per dietary condition. Vsd values from the above-mentioned list (of 55) genes were clustered using the “Pheatmap.R” package, method “complete”. Distances between samples were calculated by correlation, as described in the “pheatmap.R” package. Gene ontology analysis on all DE genes (adjusted p-value<0.1) per dietary condition was done using the “GOSeq.R” package in R^116^. P values were adjusted using p.adj in R, method Benjamini-hoch, we used the adjusted p-value<0.05 as the threshold to determine significantly regulated GO terms. For clustering based on GO terms we focused on overrepresented GO terms and used -log10(adjusted p-values) of all terms that were found significant in at least one diet (adj. p-value<0.05). Euclidean distance was used to measure distance between conditions and the clustering method used was “complete”.

### Image acquisition and analysis

For visualization of expression patterns, males from each *Gal4* line were crossed to females homozygous for the *UAS-CD8:GFP* reporter. Mated female flies with both the *Gal4* and the UAS reporter were kept on the indicated holidic medium for 3 days before dissections, unless otherwise stated. Samples were dissected in 4°C PBS, then transferred to a formaldehyde solution (4% paraformaldehyde in PBS + 10 % Triton-X) and incubated for 20–30 min at room temperature. Samples were washed three times in PBST (0.5 % Triton-X in PBS) and then mounted in Vectashield with DAPI (H-1200, Vector Laboratories). Olfactory lobe images were captured on a Zeiss LSM 980 inverted confocal microscope using a 20x objective, and Fiji ^117^ was used for imaging analysis. For image analysis, each image ROI was selected based on max-projection to cover the entire detected glomeruli. The mean fluorescent signal for the sum of intensities of the z-stack was measured within the ROI and in the background. Mean intensity was normalized by subtraction of background mean (ROI mean - Background mean).

### flyPAD assays

Food choice experiments were done using flyPAD as described in^118^. Single flies in different dietary conditions were tested in arenas containing two types of food patches: 10 % yeast (Saf-instant, Lesaffre) or 20 mM sucrose (#84097, Sigma), each mixed with 1 % agarose (#16500, Invitrogen). For all experiments, flies were individually transferred to flyPAD arenas by mouth aspiration and allowed to feed for 1 h at 25°C, 70 % relative humidity. The total number of sips per animal was calculated using previously described flyPAD algorithms^118^. Non-eating flies (defined as having fewer than two activity bouts during the assay) were excluded from the analysis. flyPADs were run in parallel to sample collection for RNA-seq, obtaining n >16 flies per condition in each repetition (“biological replicate”). Data from all replicates (n=6-7) was pooled (approximately 120 flies per dietary condition) and boot-strapping analysis was performed by sampling groups of 30 flies per dietary condition 10,000 times to evaluate population median and 95% confidence interval per dietary condition.

### Video-tracking assay

Video tracking foraging assays were performed based on previously described protocols^19,84,86^. In brief, flies were individually transferred to circular arenas by mouth aspiration and allowed to freely forage for one hour at 25°C, 70 % RH, while being video-recorded. The foraging assay was performed in an arena containing twelve 5 µL food patches, half containing 10% yeast and half 100 mM sucrose (#84097, Sigma), or each half containing a different yeast genotype, distributed in a radial pattern. The Bonsai framework was used to acquire the data^119^, and analysis was done in Python.

### Yeast for tracking assay with mutated yeast

The parental strain *Saccharomyces cerevisiae* BY4743 (MATa/α, his3ll1/his3ll1, leu2ll0/leu2ll0, met15ll0/MET15, LYS2/lys2D0, ura3ll0/ura3ll0) and the deletion mutant *Ilv2* were obtained from EUROSCARF (Frankfurt, Germany). Strains were routinely maintained in YPD medium containing 2 % (w/v) glucose (#G7528, Sigma), 2 % (w/v) yeast extract (LP0021, OXOID), 1 % (w/v) peptone (#211677, BD-Difco) and 2 % agar (#214530, BD-Difco). To prepare the yeast suspensions used in the tracking assays, cells were batch cultured in MM4 medium containing 1.7 g L^-1^ Yeast Nitrogen Base (YNB) w/o amino acids or ammonium (#233520, DB-Difco), 20 g L^-1^ glucose, 2.65 g L^-1^ (NH_4_)_2_SO_4_ (#09978, Sigma-Aldrich), supplemented with 20 mg L^-1^ methionine (#M9625, Sigma), 30 mg L^-1^ lysine (#L5626, Sigma), 60 mg L^-1^ leucine (#L8912, Sigma), 20 mg L^-1^ histidine (#H8000, Sigma) and 20 mg L^-1^ uracil (#U1128, Sigma). An inoculum with 500 ml of MM4 medium and with an initial optical density at 600nm (OD_600nm_) of 0.01 were prepared for each strain. The cells were cultivated at 30°C and with orbital agitation at 180 rpm. On the next day, when an OD_600nm_ of 1.0 +/-0.2 was reached, the cells were spun down by centrifugation at 3000xg for 10 min, washed with 10 ml of sterile water and centrifuged again to collect the cells. The pellet was then resuspended in 2 ml of sterile water, 20 mg of agarose (#16500, Invitrogen) was added and the cell suspension. The mixture was heated at 70°C for 30 min and used as a food source in the tracking assays.

### Preparation of bacterial suspensions for tracking assays

To prepare the *Lactobacillus brevis*^EW 120^ suspension, 40 ml of liquid MRS broth (Fluka, #38944) was inoculated with bacteria and incubated overnight at 37°C without orbital shaking. To prepare the suspensions with 150 OD/ml used in the tracking assays, the OD was measured, and the volume of culture necessary to prepare the final volume of suspension to be used in the tracking assay was calculated. Then, the volume of bacteria culture calculated was centrifuged and the supernatant was discarded. Finally, the pellet was resuspended in fresh MRS at the volume appropriate to prepare the desired cellular suspension at 150 OD/ml. The bacterial suspensions were always prepared fresh on the day of the tracking assay.

### *In-vivo* Calcium imaging

Seven to eleven-day-old female flies were immobilized with their eyes and thorax secured below a metal plate using fast-curing UV glue (Bondic). A slit in the metal allowed access to the head from above. For visual access to both antennal lobes, a window was cut in the head capsule between the antennae and ocelli. The head was covered with hemolymph-like saline solution (108 mM NaCl, 5 mM Kcl, 4 mM NaHCO_3_, 1 mM NaH_2_PO_4_, 5 mM Trehalose(D+) dihydrate, 5 mM HEPES, 10 mM Sucrose, 2 mM CaCl_2_ 2H_2_O, 8.2 mM MgCl_2_ 6H_2_O).

Imaging was performed on a resonant-scanning two-photon microscope (Scientifica) using a 20x NA 1.0 water immersion objective (Olympus), controlled by a piezo-electric z-focus drive. Volumes covering glomeruli on both hemispheres were obtained at 512 × 256 × 16 voxels with a zoom level of 1.5x and a volume rate of ∼ 4Hz.

Odor stimulation was performed using a custom-made olfactometer^121^ 20 µl odor dilutions were pipetted onto syringe filters (Puradisc 13, Whatman). Carrier- and stimulation air flow were regulated by two flow meters (TW-32900-44, Cole Parmer). Odor valves (LHDA 1233115H, The Lee Company) were controlled using an Arduino Uno (Arduino) to direct airflow over the different odor channels to inject odorized air into a carrier airstream. Airstreams were combined using a custom-made polyetheretherketone manifold. Per odor, we performed two consecutive 1s stimulations, spaced by 5 s.

### Ca^2+^ imaging data analysis

Data pre-processing was performed in Python. A Gaussian filter of 3×3 pixels was applied to the maximum intensity projections of the recorded volumes. Data was then corrected for movement in x and y using the *MotionCorrect* function provided by CaImAn^122^. For each fly, regions of interest (ROIs) were defined based on the maximum intensity projection for this fly, then processed by using the threshold tool followed by particle analysis, with the lowest size of 100^2^ pixels. Firstly, the mean fluorescence intensity within the ROI at each time point. Secondly, each trial was then aligned to stimulus onset based on the change in signal >2 compared to baseline (baseline=mean fluorescence at the beginning of each trial). For each trial and ROI, the mean fluorescence intensity within the ROI at each time point was used as *F*, and *F*_*0*_ was calculated as the median of these values from 5 seconds before the detected stimulus onset. ll*F/F*_*0*_ was re-calculated as (*F-F*_*0*_)/*F*_*0*_ for each time point. To calculate the area under the curve, we took the sum of the ll*F/F*_*0*_ values from 2 to 44 seconds after the first stimulus onset for each fly’s normalized fluorescence.

## Odorant dilutions

Odor stimuli were prepared in MilliQ water (Merck). Phenethylamine was used at vol/vol dilutions of 10^−6^.

## Statistics

For all the plots statistic tests indicated in the figure legend were done using R^123^.

## Additional resources

Holidic diet recipe: http://dx.doi.org/10.17504/protocols.io.heub3ew

## Code

This paper does not report original code.

## Author Contributions

Conceptualization, G. E. N. and C.R.; Methodology, G.E.N., S.H., D.M., C.R.; Validation, G.E.N., S.H., D.M.; Data Curation and formal analysis, G. E. N. and C.R.; Investigation, G. E. N., S.H., D.M., A.P.F., C.B, A.P.E.; Supervision, C.R.; Data curation, G.E.N; Visualizations, G.E.N., S.H., C.R.; Writing – Original Draft, G. E.N. and C.R.; Writing – Review & Editing, G.E.N., S.H., D.M., B.D., A.P.E, A.P.F., C.B., C.R.; Project Administration G.E., C.R.; Funding acquisition, G.E., C.R.. All authors read and approved the final manuscript.

