## Supplemental figures 1-3 for "Lack of Single Amino Acids Transcriptionally Remodels Sensory Systems to Enhance the Intake of Protein and Microbiota"

### Supplementary figures

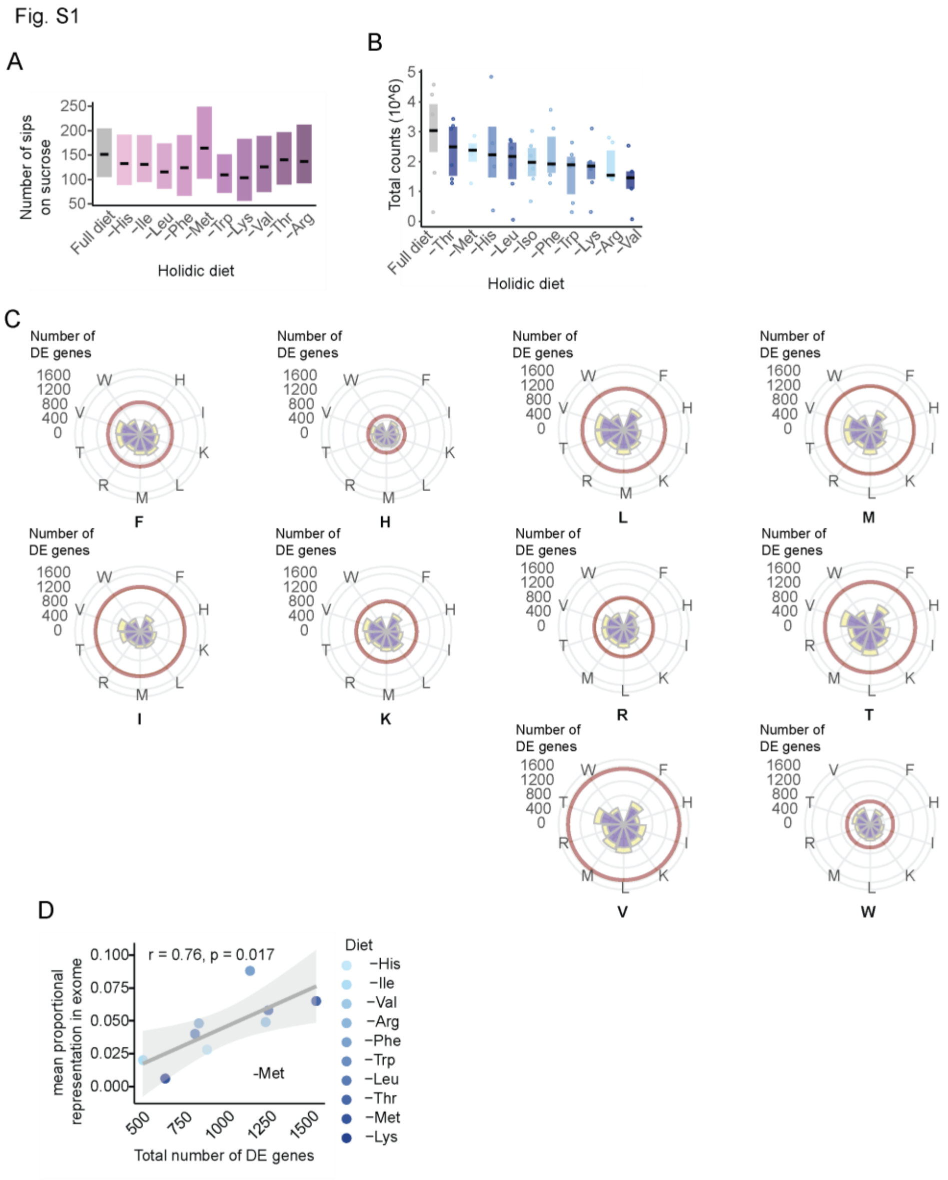


**Fig. S1. Single amino acid deprivation induces changes in head transcriptome but does not affect sucrose feeding**

**(A)** Black lines represent the medians and the boxes represent 95% confidence interval (CI) of number of sips on sucrose of flies deprived of each of the 10 eAAs. **(B)** Total number of sequenced counts per sample and dietary manipulation. Black lines represent median values. Boxes represent IQR. Each datapoint represents a pooled sample of 10 fly heads. n=5-7. **(C)** Circular histogram depicting the number of shared significantly upregulated (yellow) and downregulated (purple) genes amongst the different deprivation conditions. Circles depict the number of genes as indicated in the legend on the left of the plot. The red circle indicates total number of significantly regulated genes (adjusted p-value<0.1) of the eAA stated below each histogram. **(D)** Pearson correlation between the number of DE genes, excluding Met, and their relative representation in the exome (according to Piper et al., 2017). Each datapoint represents the median value corresponding to a single eAA deprivation from Fig. 1C and their matching unique differentially expressed gene numbers or representation in the exome. Shading represents 95% CI.

Fig. S2

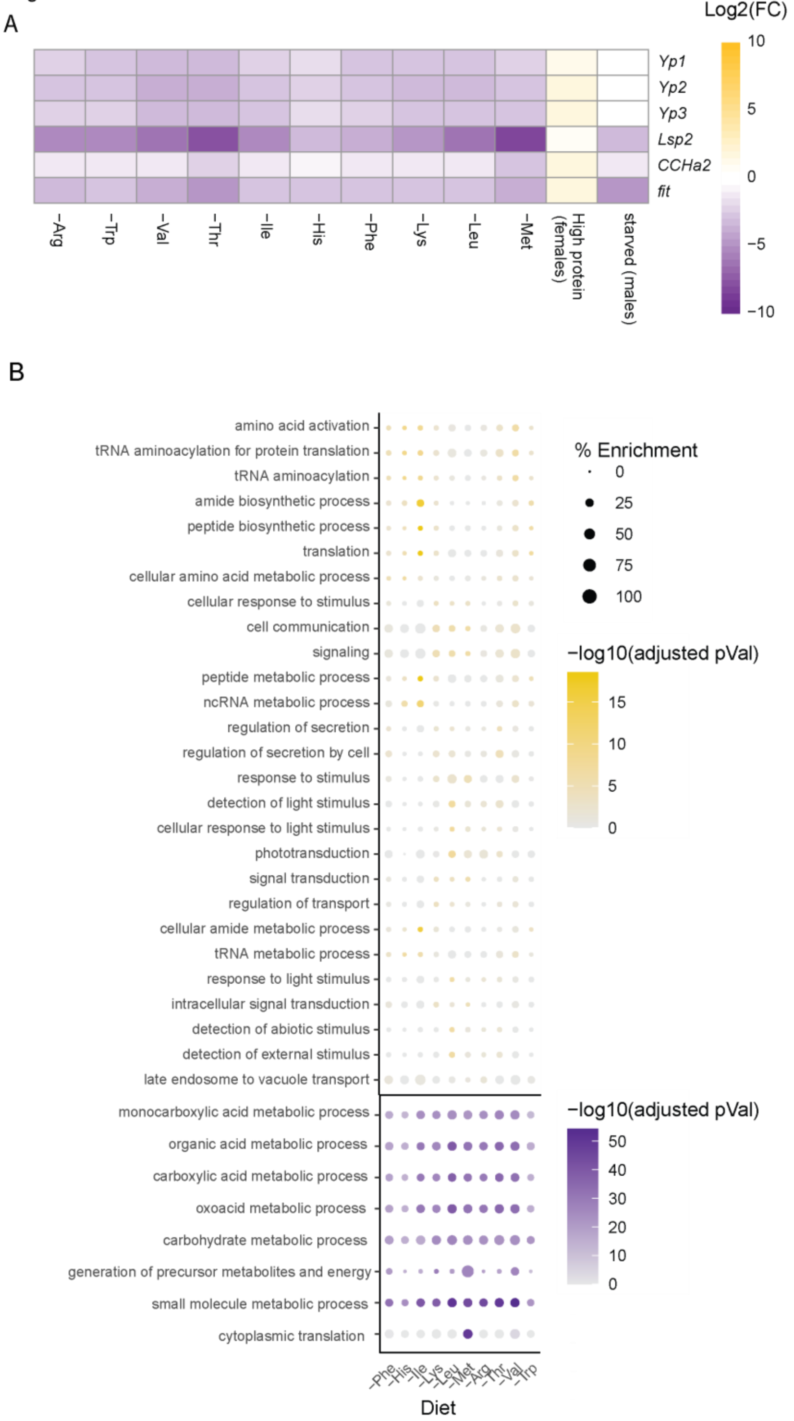


**Fig. S2. Ile and Met deprivations induce a unique transcriptional regulation fingerprint in the fly head related to transcription and translation GO terms, respectively.**

**(A)** heatmap depicting log2(FC) values of the selected established metabolic genes in our dataset and compared to published data of female flies maintained on high yeast diet [78] and starved adult males [79]. **(B)** Most significantly over represented gene ontology terms identified based on upregulated genes (yellow) or downregulated genes (purple) for each eAA depletion (top 20 per eAA). Circle size (% enrichment) was calculated by dividing number of differentially expressed genes within a category by total number of genes in a category. Color intensity represents FDR adjusted p-values.

Fig. S3

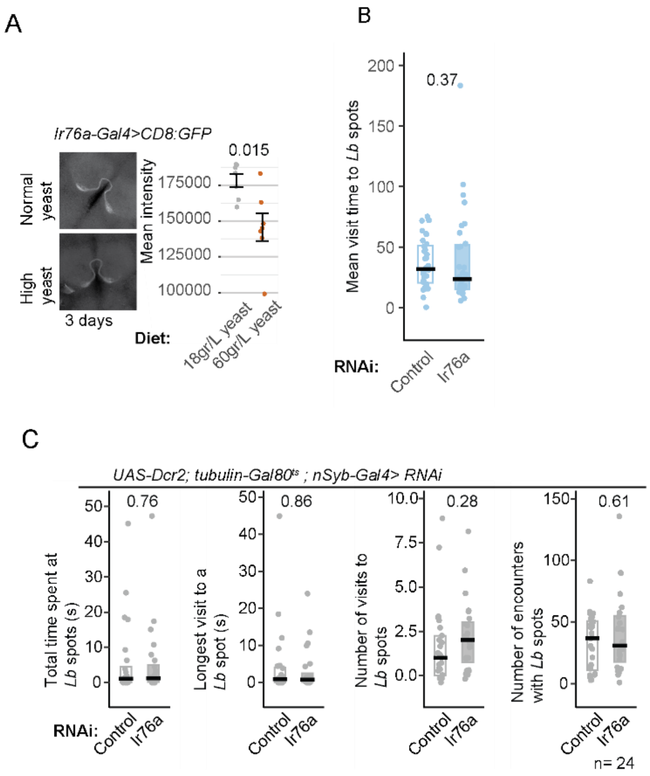


**Fig. S3. *Ir76a* levels correlate with protein and AA levels in the diet and with behavioral approach to lactobacilli**

(**A)** Reporter based analysis of *Ir76a* regulation by yeast content in the diet. Left panel: representative sum of projections confocal images of antennal lobes obtained from *Ir76a-Gal4> CD8:UAS-GFP* flies following 3 days on food containing 18gr yeast/L or 60gr yeast/L. Right panel: quantification of normalized mean fluorescence intensity in *Ir76a* glomeruli. Data presented as +- SEM. Each datapoint corresponds to an individual fly. Numbers at the top of the plots indicate p values as calculated using the t-test; n=7. **(B)** Mean duration of *Lb* patch visits of Ile deprived flies expressing RNAi against *Ir76a* (full light blue) or control RNAi (open light blue) driven by *UAS-Dcr2; tubulin-Gal80$^{ts}$; nSyb-Gal4*. Black line represents medians; Boxes represent IQR; Each datapoint corresponds to a single fly. Numbers at the top of the plots indicate p values as calculated using the Wilcoxon signed rank test, n=28. **(C)** *Lb*-tracking assay results of fully-fed flies expressing RNAi against *Ir76a* (full gray boxes) or control RNAi (open gray boxes) driven by *UAS-Dcr2; tubulin-Gal80$^{ts}$; nSyb-Gal4;* Driver line indicated at the top of all plots. Black line represents medians. Boxes represent IQR. Each data point corresponds to a single fly. Numbers at the top of the plots indicate p values as calculated using the Wilcoxon signed rank test. n=24.
